# Interpretable deep learning approach for extracting cognitive features from hand-drawn images of intersecting pentagons in older adults

**DOI:** 10.1101/2023.04.18.537358

**Authors:** Shinya Tasaki, Namhee Kim, Tim Truty, Ada Zhang, Aron S Buchman, Melissa Lamar, David A. Bennett

## Abstract

Hand drawing involves multiple neural systems for planning and precise control of sequential movements, making it a valuable cognitive test for older adults. However, conventional visual assessment of drawings may not capture intricate nuances that could help track cognitive states. To address this issue, we utilized a deep-learning model, PentaMind, to examine cognition-related features from hand-drawn images of intersecting pentagons. PentaMind, trained on 13,777 images from 3,111 participants in three aging cohorts, explained 23.3% of the variance in global cognitive scores, a comprehensive hour-long cognitive battery. The model’s performance, which was 1.92 times more accurate than conventional visual assessment, significantly improved the detection of cognitive decline. The improvement in accuracy was due to capturing additional drawing features that we found to be associated with motor impairments and cerebrovascular pathologies. By systematically modifying the input images, we discovered several important drawing attributes for cognition, including line waviness. Our results demonstrate that hand-drawn images can provide rich cognitive information, enabling rapid assessment of cognitive decline and suggesting potential clinical implications in dementia.

## Introduction

Copying geometric or abstract figures is a complex behavior that requires the integration of multiple cognitive domains, including executive function to initiate the task, visuospatial abilities to carry it out, and, to a lesser extent, semantic memory to produce the correct image. Therefore, paper-and-pencil drawing tasks are often employed independently or as part of a larger screening tool to detect cognitive impairment, including Alzheimer’s dementia^1,2^, and Parkinson’s disease^3,4^. For example, copying intersecting pentagons, the Pentagon Drawing Test (PDT), is used as one of the items in the 30-item Mini-Mental State Examination (MMSE) - a common tool to evaluate a person’s mental health and identify potential cognitive impairments^5^. PDT involves asking the participants to draw two intersecting pentagons on a piece of paper. In the MMSE, the intersecting pentagons are rated simply 0 (fail) or 1 (pass) based on limited factors. More detailed evaluations of the intersecting pentagons have been shown to provide granular-level information about a person’s cognitive abilities and potential dementia status^1,6^.

While paper-and-pencil drawing tests, like the PDT, can be a useful tool in assessing a person’s cognitive abilities, the conventional scoring has some limitations. First, these tests are often prone to subjective assessment and rater bias with different raters potentially having different interpretations of the drawings, which can lead to inconsistencies in the scoring and potentially impact the accuracy of the results. Secondly, these tests can be labor-intensive and time-consuming to score, particularly when used within a larger population. Finally, these tests are typically focused on a limited set of drawing attributes. For example, the PDT scoring is based on only a few factors, such as the presence or absence of intersections and the number of vertices^5^, which only capture limited aspects of behavior and hence cognitive status^1^. Thus, while paper-and-pencil drawing tests can be useful, their human-based scoring has limitations in scope, accuracy and scalability.

Automated scoring methods that utilize machine learning methods offer the potential to address some or all of these weaknesses. For example, deep-learning techniques have demonstrated promising performance in automating ratings for the PDT^7,8^, Rey Complex Figure Test^9,10^, and Clock Drawing Test^11,12^. However, the primary objective of the previous automated scoring is to reproduce human-based conventional ratings and few machine learning approaches directly predict cognitive performance from drawings^13^. In addition, prior work has not been able to explore the factors that may confound or mediate the characteristics of cognitive impairment in intersecting pentagons, such as motor abilities and brain pathologies. Generating such integrated mechanisms requires drawing images, various phenotypes, and multiple brain pathologies assayed in the same set of individuals and the analytical frameworks to integrate these data. Furthermore, none of the studies utilized the models to discover the novel drawing features associated with cognitive impairment, which may be important for advancing our understanding of visuospatial memory ability and motor coordination of drawing in older adults.

Here, we leveraged data from 3,111 participants from three ongoing cohort studies of aging and dementia at the Rush Alzheimer’s Disease Center (RADC) to train a deep learning model that predicts global cognition performance (Figure 1A). By performing a thorough evaluation of 47 established deep-learning models for vision recognition, we identified an architecture that demonstrated high and robust performance. Along with detailed cognitive tests, we were also able to incorporate comprehensive motor examinations conducted with participants and a variety of neuropathologies recorded for several hundred of participants. We used these data to break down the non-cognitive and pathological components of the model’s prediction. Furthermore, to address the challenge of interpretability of deep learning models, we developed a pentagon-drawing simulator. This simulator is an explainable-AI approach that allowed us to interrogate the deep learning model to suggest the key drawing characteristics in people with lower cognitive function.

**Figure 1.**
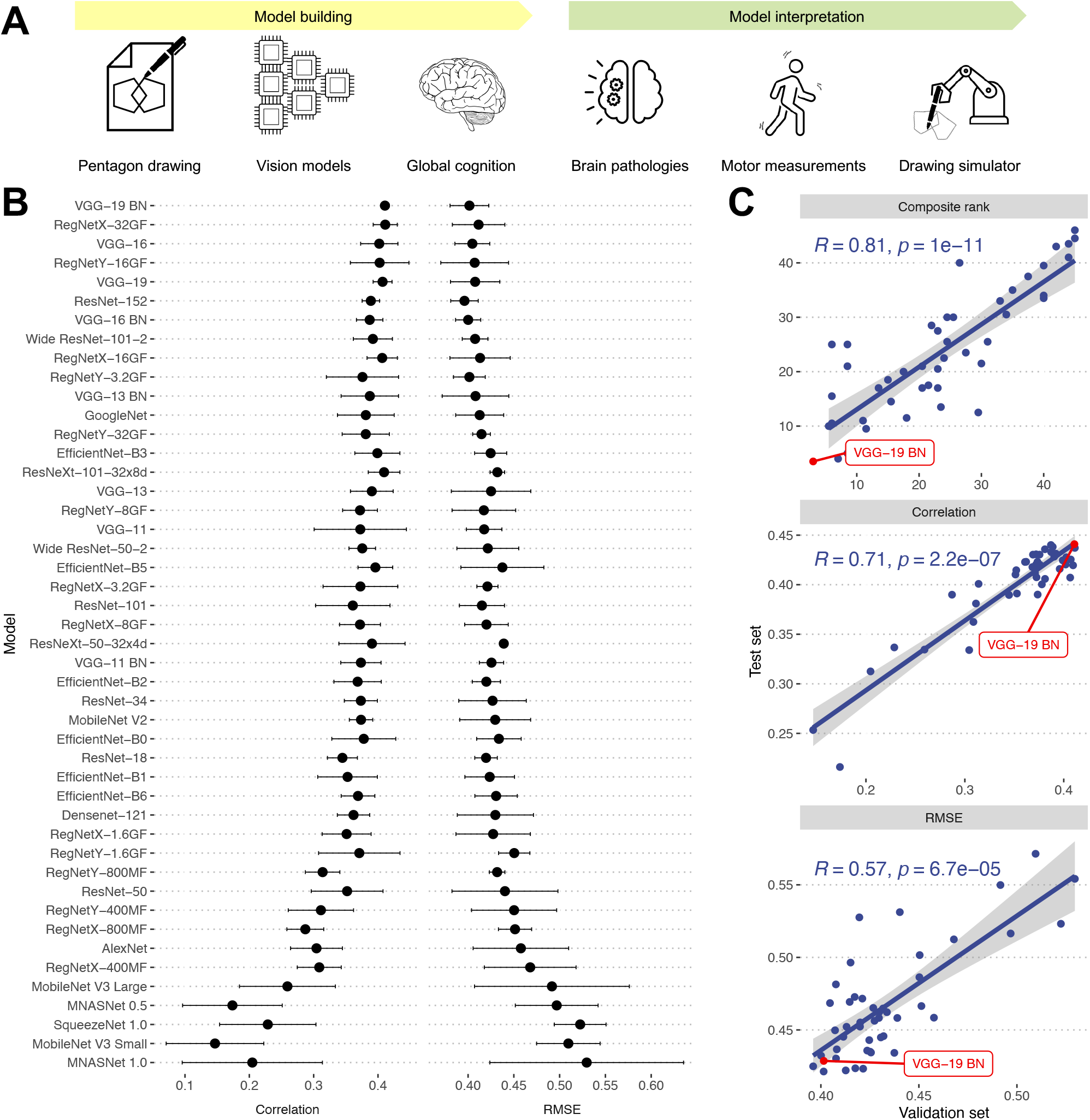
Development of a deep learning model for predicting global cognitive performance. (A) Schematic representation of the model development and interpretation process for predicting global cognition performance. The process includes (1) Model Building: selection of a high-performing vision recognition architecture from 47 established deep-learning models, and (2) Model Interpretation: integration of comprehensive motor examinations, neuropathologies, and development of a pentagon-drawing simulator to identify key drawing characteristics in individuals with lower cognitive function. (B) Validation performances of the 47 deep learning models. Spearman’s correlation and RMSE between predicted cognition scores from each model and measured values were computed for validation samples. We repeated model training five times, each time using a distinct training set. The Median and median absolute deviation of the metrics from the five runs was plotted. Models were ranked based on the average ranking of Spearman’s correlation and RMSE. (C) Comparison of model’s performances between validation and test sets. The composite ranking was obtained as an average of rankings based on Spearman’s correlation and RMSE. The composite ranking, Spearman’s correlation, and RMSE are compared between validation (x-axis) and test (y-axis) sets. The based on the VGG-19 BN architecture is highlighted as it showed the best permanence with the validation sets.

## Results

### Characteristics of study participants

Participants enrolled at the age of 77.4 (SD: 7.6) with a follow-up of 5.6 years (SD: 4.6) (Table 1), on average. Of participants, non-Latino white (77.6%) is the most common race, followed by African American (21.6%). At yearly home visits, participants received comprehensive cognitive assessments, which included 19 tests used to generate a composite measure of global cognition. In addition, the pentagon drawing test was administered as one of the items of the standard MMSE. Note that the pentagon drawing test is not included in global cognition which is based on 19 measures independent of MMSE. We scanned 13,777 drawings of intersecting pentagons obtained throughout the visits.

**Table 1.**
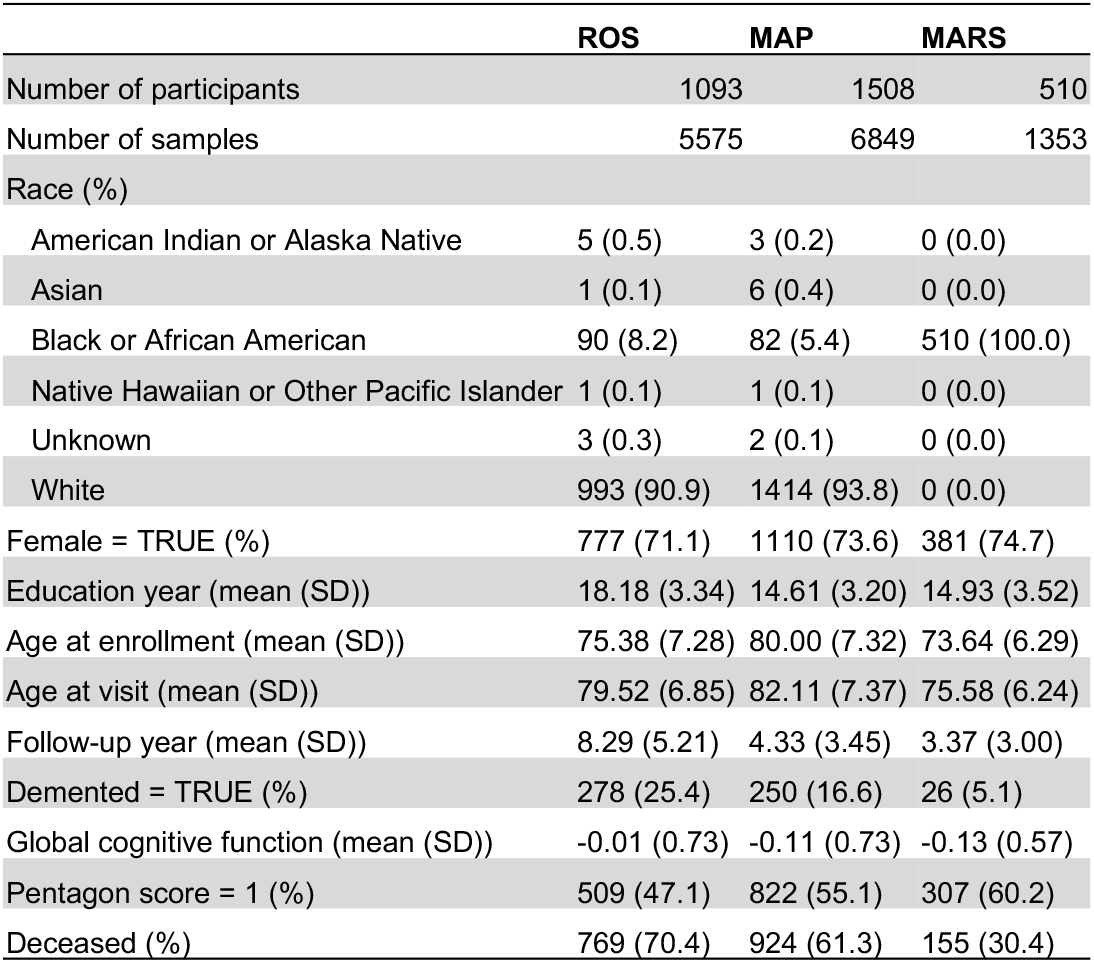
Demographic information for the ROS, MAP, and MARS cohorts.

|  | ROS | MAP | MARS |
| --- | --- | --- | --- |
| Number of participants | 1093 | 1508 | 510 |
| Number of samples | 5575 | 6849 | 1353 |
| Race (%) |  |  |  |
| American Indian or Alaska Native | 5 (0.5) | 3 (0.2) | 0 (0.0) |
| Asian | 1 (0.1) | 6 (0.4) | 0 (0.0) |
| Black or African American | 90 (8.2) | 82 (5.4) | 510 (100.0) |
| Native Hawaiian or Other Pacific Islander | 1 (0.1) | 1 (0.1) | 0 (0.0) |
| Unknown | 3 (0.3) | 2 (0.1) | 0 (0.0) |
| White | 993 (90.9) | 1414 (93.8) | 0 (0.0) |
| Female = TRUE (%) | 777 (71.1) | 1110 (73.6) | 381 (74.7) |
| Education year (mean (SD)) | 18.18 (3.34) | 14.61 (3.20) | 14.93 (3.52) |
| Age at enrollment (mean (SD)) | 75.38 (7.28) | 80.00 (7.32) | 73.64 (6.29) |
| Age at visit (mean (SD)) | 79.52 (6.85) | 82.11 (7.37) | 75.58 (6.24) |
| Follow-up year (mean (SD)) | 8.29 (5.21) | 4.33 (3.45) | 3.37 (3.00) |
| Demented = TRUE (%) | 278 (25.4) | 250 (16.6) | 26 (5.1) |
| Global cognitive function (mean (SD)) | -0.01 (0.73) | -0.11 (0.73) | -0.13 (0.57) |
| Pentagon score = 1 (%) | 509 (47.1) | 822 (55.1) | 307 (60.2) |
| Deceased (%) | 769 (70.4) | 924 (61.3) | 155 (30.4) |

### Model architecture for predicting cognition via drawing (PentaMind)

To develop models that predict global cognition from intersecting pentagon drawings, we comprehensively evaluated 47 published convolutional neural network models for image recognition (Methods). The 47 models were designed and trained to categorize images into 1,000 object classes^14^. We modified the model structures so that the final output layer outputs a numeric value instead of object classes. As we collected multiple images from the same participant across the period, we used a person-based split rather than an image-based split to ensure that models were not trained with images from the same participants in both the training and validation/test sets. Specifically, we randomly divided 3,111 participants into a training set (80%), a validation set (10%), and a test set (10%). To improve the reliability of performance evaluation, we repeated the training procedure five times using five distinct sets of training and validation samples.

Figure 1B indicates the comparative performance of the models based on Spearman’s correlation and root mean squared error (RMSE) for the validation sets. We selected a model that demonstrated superior performance in both correlation and RMSE metrics for further investigation. Specifically, we calculated a composite ranking as an average of the rankings based on Spearman’s correlation and RMSE. The model that ranked highest overall among the 47 models was the model based on VGG19 with batch normalization (VGG19-BN), which we named PentaMind. This model ranked second in Spearman’s correlation (0.41) and fourth in RMSE (0.40). The accuracy of PentaMind on the test sets was reasonably high, with a Spearman’s correlation of 0.44 and an RMSE of 0.42. Importantly, the performances of models on validation and testing sets are highly congruent, indicating that selecting a representative model based on validation sets did not introduce bias (Figure 1C).

### Evaluation of PentaMind using test dataset

To conduct a series of evaluations for our PentaMind, we predicted the global cognition from all 13,777 images. However, applying the model to training samples may introduce bias into the estimates. To prevent this, we retrained PentaMind using an out-of-fold prediction strategy. Specifically, we divided the images into ten non-overlapping folds.

Then, we trained the model using nine folds and tested on the left-out tenth fold. This procedure was repeated for each of the ten folds, resulting in ten models, each trained on different sets of training and holdout images. By concatenating the predicted global cognitive function for the holdout samples, we obtained an unbiased prediction for all 13,777 images, which could then be used to calculate overall prediction accuracy and examine the characteristics of our prediction (Figure S1).

The predicted cognition score explained 23.3% of the variance in global cognition, which is 1.92 times greater than that of manual standard binary scoring (Figure 2A and Table S1) that is based on the presence or absence of intersections and the number of vertices. A regression model using PentaMind-predicted scores and human binary scores as covariates was able to explain 24.7% of the variance in global cognition together. Notably, the PentaMind’s predictions were the primary contributor as compared to the binary scoring, independently explaining 18.0% of the variance (Table S1). Next, we investigated whether PentaMind could differentiate cognitive status with the images scored as 1 (pass/normal) by the conventional clinical evaluation. Intriguingly, the PentaMind exhibited a Spearman correlation of 0.32, accounting for 10.0% of the variance in the global cognition score (Figure 2B and Table S2). The PentaMind accounted for more variance (24%) in a group of failed pentagons (score = 0 from binary scoring), but there was no significant difference in variance explained between the groups (p=0.41). These findings demonstrate that our PentaMind can capture nuanced characteristics of human drawings that are undetectable, unquantifiable, or omitted by the conventional clinical assessment.

**Figure 2.**
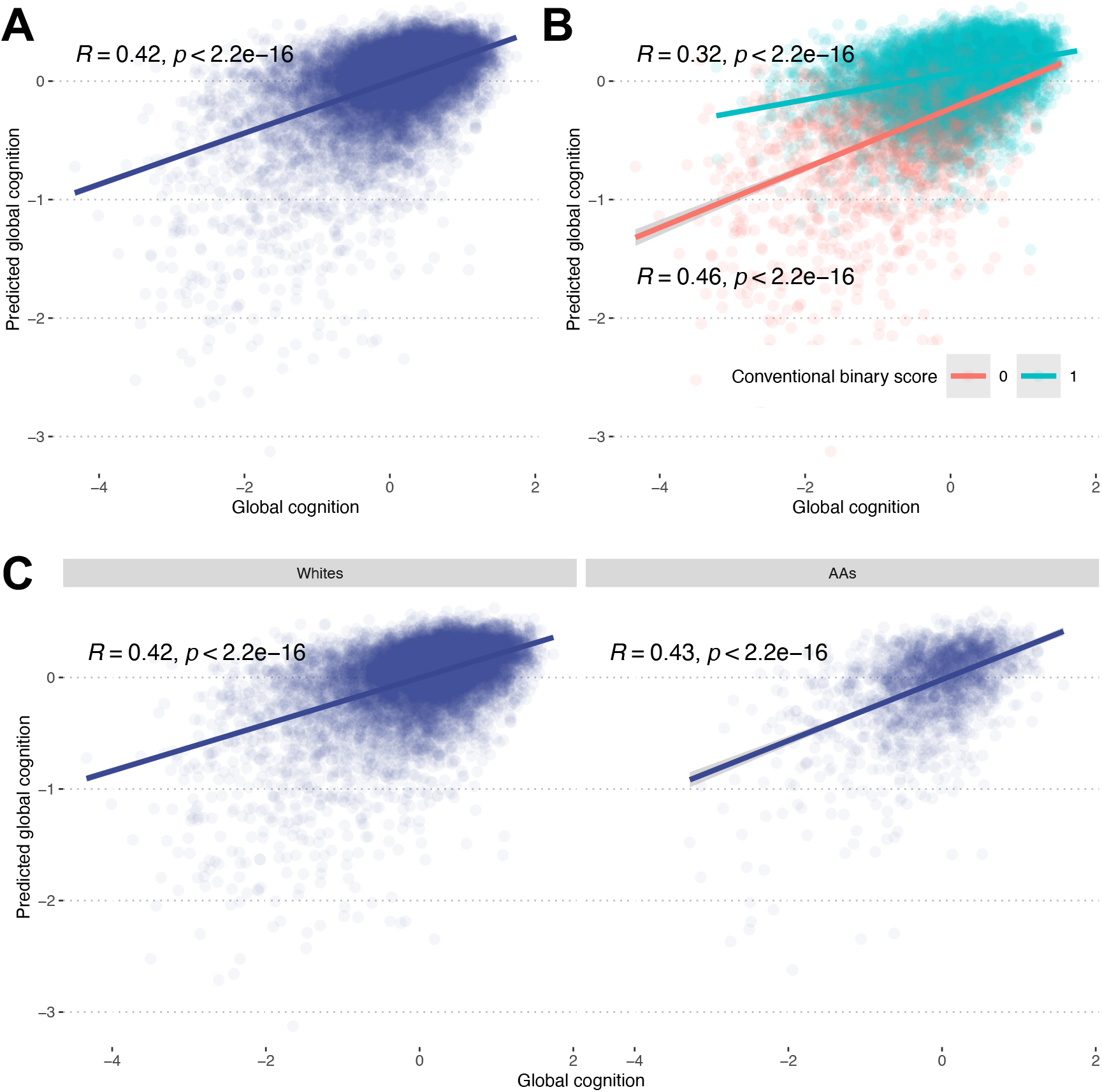
Performance evaluation of PentaMind. (A) Relationship of the predicted cognition scores and actual scores. We employed an out-of-fold prediction strategy to generate an unbiased predicted cognition score for all 13,777 images. The scatter plot compares the actual and predicted cognition scores with a linear regression line. Spearman’s correlation and p-value are also displayed. (B) PentaMind’s performance stratified by the conventional binary rating. The images are stratified based on the conventional rating: success (1) or failure (0). Spearman’s correlation and p-value for each group are displayed. (C) PentaMind’s performance stratified by the race of participants. The performance metric is calculated for Whites and African Americans separately.

### Generalization of PentaMind across race

By leveraging the racial diversity of our observational community-based cohorts, we examined the generalizability of our model across race. To do this, we calculated prediction performance separately for white participants (77.6% of participants) and those identifying as African American (21.6% of participants). The predictive performance based on Spearman’s correlation for the two racial groups were 0.42 and 0.43, respectively (Figure 2C), while the percentage of variance in cognitive score explained by PentaMind for white and African American participants were 23.2% and 24.1%, respectively (Table S3). This comparable performance demonstrates that our model may successfully generalize across both white and African American participants.

### Relationship of PentaMind’s prediction with clinical phenotypes

Global cognition is a summary representation of various aspects of cognitive functions. To understand whether the improvement of PentaMind over the conventional clinical assessment is attributed to the model’s ability to capture specific cognitive components, we analyzed various phenotypic measurements acquired simultaneously from the same individual. Specifically, we explored the link of the predicted global cognition with five domains of global cognition and ten motor functions that are known to be associated with cognitive impairment (Table S4). Motor function is a complicated action that may necessitate the use of many clinical instruments to capture the various deficiencies that appear in older adults. Therefore, we investigated two interrelated motor phenotypes, a global parkinsonism score and a global motor score, and their domains.

All 15 phenotypes were significantly associated with global cognition score predicted by PentaMind (Bonferroni-corrected-p < 0.05), but only eight were associated with the conventional binary scoring (Figure 3 and Table S5). Overall, PentaMind accounted for about five times more variance than the manual binary scoring alone. Notably, this tendency was more pronounced for motor-related phenotypes than for cognition domains, where the predicted score of PentMind was significantly more strongly associated with motor phenotypes than the manual binary scores, with a magnitude of 6.4 times. While for cognitive scores, the predicted score was also more strongly associated than the manual binary scores, the effect size was smaller, with a magnitude of 2.6 times. The result implies that PentaMind’s improved performance is partly due to its capability to extract signals pertaining to motor impairment from a handwriting image that may be associated with global cognition. The result also highlights the limitation of the conventional scoring, which is dependent on basic indicators such as the number of vertices and the presence of the intersection of two pentagons.

**Figure 3.**
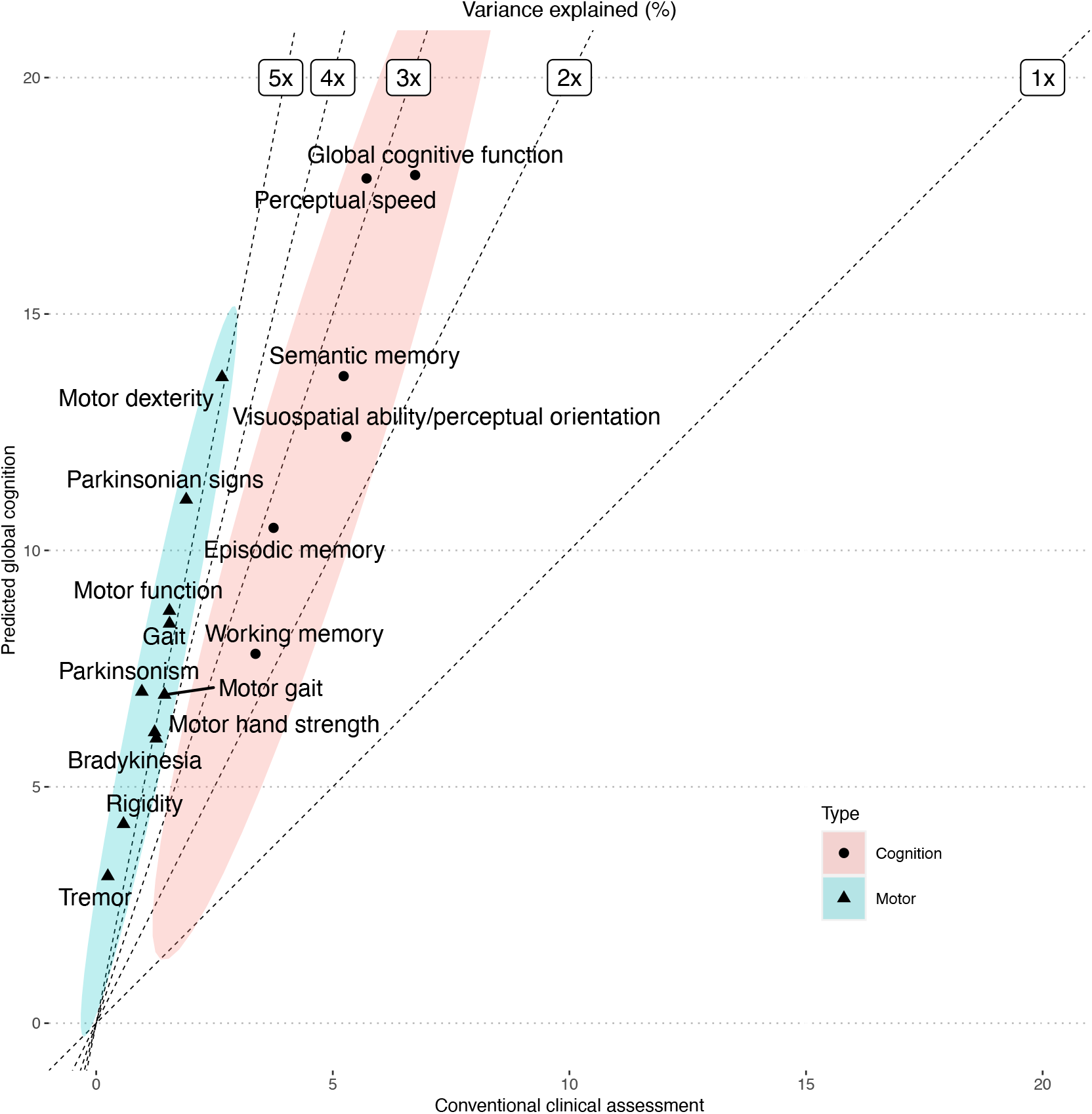
Relationship of Pentamind’s prediction with clinical phenotypes. The percent of the variance in each clinical phenotype explained by the predicted cognition (y-axis), and the conventional clinical rating of PDT (x-axis) is compared. Dashed lines indicate the fold improvements in percent of variance explained by the PentaMind over the conventional rating.

### Relationship of PentaMind’s prediction with brain pathologies

The existence of brain pathologies is the leading cause of cognitive impairment. Consequently, it is important to determine if the global cognition score predicted by the pentagon represents cognitive impairment tied to specific brain pathologies. Therefore, we associated a broad spectrum of brain pathologies, including classical AD pathologies, Lewy bodies, TDP-43, and cerebrovascular pathologies, with the global cognition score predicted by PentaMind based on the PDT closest to death (Table S6). Out of 20 pathologic indices, nine were associated with the predicted global cognition (Bonferroni-corrected-p < 0.05), while six were associated with the conventional binary score (Figure 4 and Table S7). Comparable effect sizes for classical AD pathologies, including NIA-Reagan and global AD pathology, were observed between the predicted global cognition by PentaMind and the conventional score. By contrast, the PentaMind’s prediction showed stronger associations with cerebrovascular disease, specifically vessels disease of atherosclerosis and arteriolosclerosis, as well as loss of pigmented neurons in the substantia nigra.

**Figure 4.**
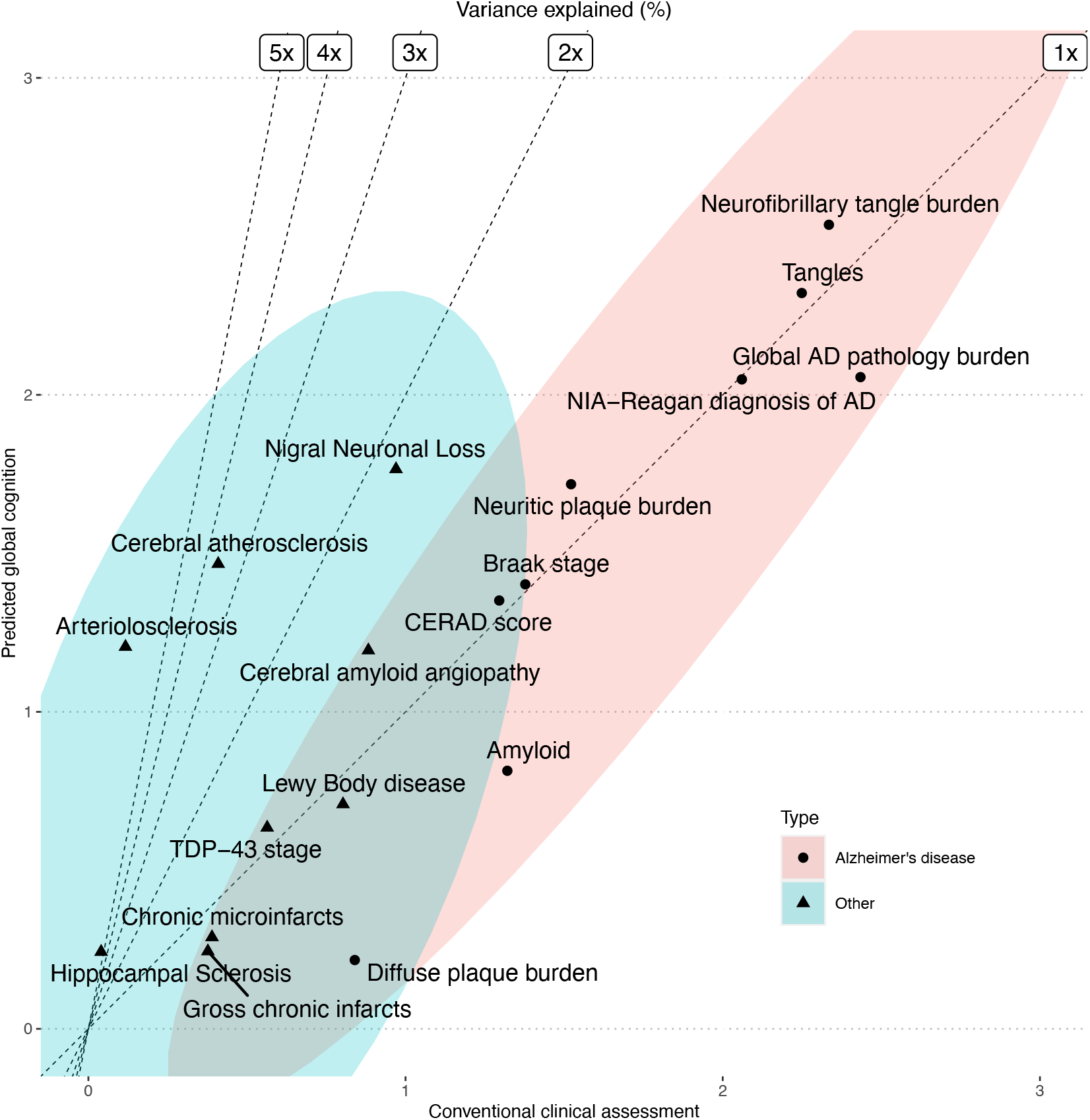
Relationship of Pentamind’s prediction with brain pathologies. The percent of the variance in each brain pathology explained by the predicted cognition and the conventional clinical rating of PDT is compared. Dashed lines indicate the fold improvements in percent of variance explained by the PentaMind over the conventional rating.

Since PentaMind appeared to have an advantage in detecting motor-related cognitive impairment as outlined above, these pathologies might be tied to their known effects on motor function^15^. To examine this hypothesis, we contrasted motor dexterity with the 20 brain pathologies. We found that the cerebrovascular pathologies of atherosclerosis and arteriolosclerosis and nigral neuronal loss were more robustly associated with motor dexterity than the other classical AD pathologies (Table S8). These congruent clinical and pathological associations indicate that the deep learning approach can detect the signs of motor dysfunction from pentagon drawing.

### Identification of crucial drawing features

Although PentaMind improved performance in predicting global cognition from the pentagon drawing, elucidating the drawing features contributing to this accuracy is crucial for gaining novel clinical insights. To pinpoint vital elements within the image, we utilized DeepSHAP^16^, a technique for comprehending how a model generates its predictions (Figure S2). DeepSHAP indicated that the model penalizes the poor shape of the pentagon and the lack of well-formed interlocking pentagons. However, it was challenging to specify key drawing features from the result of SHAP. Therefore, we developed a simulator to generate intersecting pentagons with specified parameters. By providing the simulated images to PentaMind and monitoring the predicted cognition values, we were able to identify influential drawing features linked to cognition. To generate a synthetic pentagon drawing, the simulator takes eight parameters, including (1) the number of vertices, (2) the distance between pentagons, (3) alignment of two pentagons, (4) angle distortion, (5) size equality, (6) pentagon size, (7) line width, and (8) line waviness.

First, we examined the number of vertices and the presence of intersections – two qualities that are evaluated in the conventional clinical assessment (Figure 5). As the number of vertices in a drawing increases or decreases, PentaMind’s prediction of global cognition score drops, demonstrating that the model accurately recognizes the geometry of the pentagon (Figure 5A). Additionally, the distance between two pentagons affects the predicted cognition. Consistent with the conventional evaluation, we observed a lower cognition score when no intersection was present (Figure 5B). However, PentaMind indicated that excessive overlap is also indicative of diminished cognitive ability (Figure 5B). Furthermore, the parts of pentagons that intersected influenced the prediction. Specifically, the prediction for global cognition score was highest when vertices of the two pentagons overlapped. However, the predicted cognition score was reduced when one pentagon’s vertex intersected the other’s side (Figure 5C). This is noteworthy given that in the conventional assessment, any sort of overlap is a prerequisite for pass of the PDT. Nonetheless, PentaMind model dissects the overlaps with respect to cognitive performance in greater detail. This demonstrates how PentaMind can quantify nuanced drawing features.

**Figure 5.**
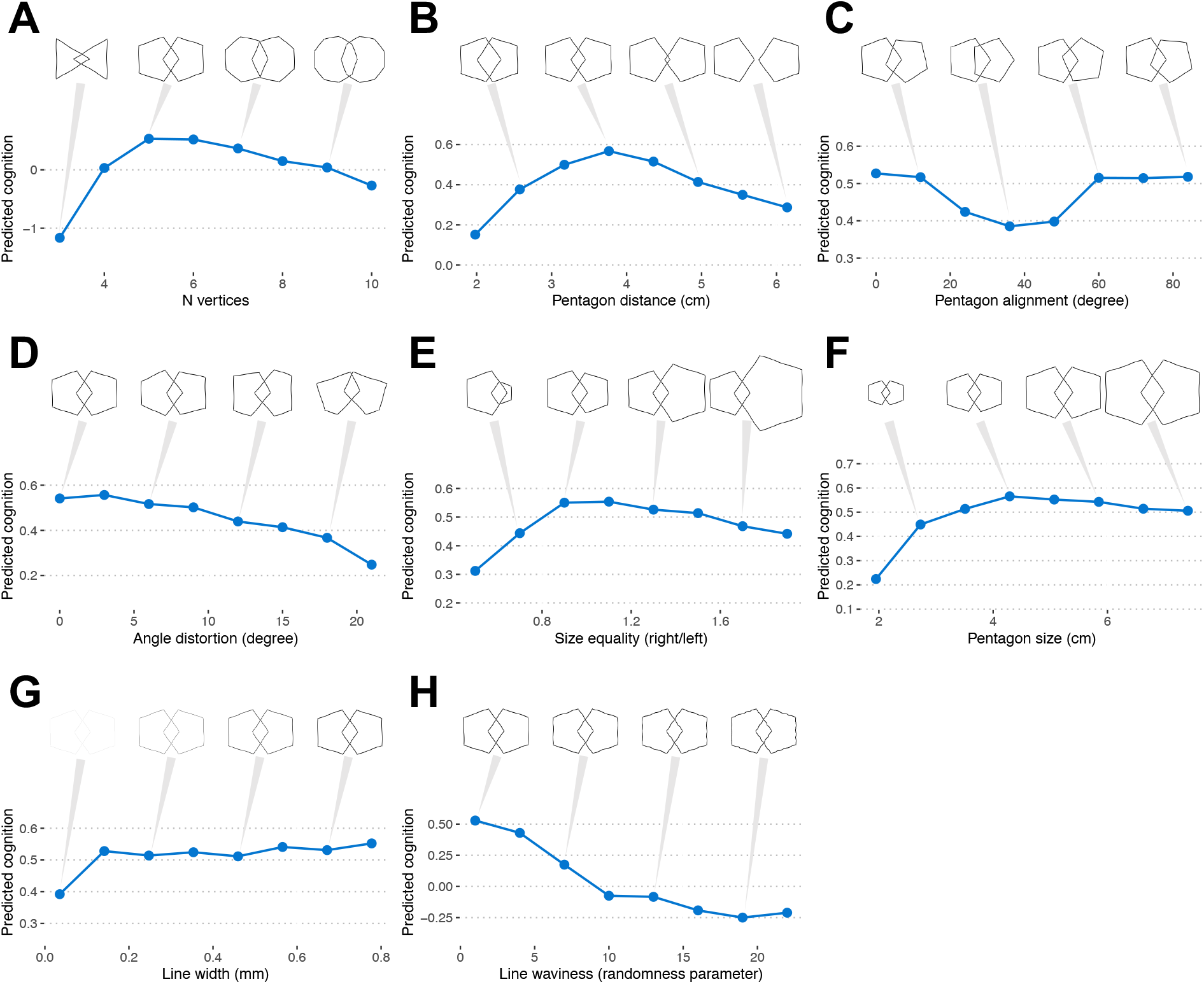
Identification of crucial drawing features. The effect of eight drawing attributes on predicted cognition sore is examined using synthetic images of PDT. The eight drawing attributes include (A) the number of vertices, (B) the distance between pentagons, (C) alignment of two pentagons, (D) angle distortion, (E) size equality, (F) pentagon size, (G) line width, and (H) line waviness. A point represents the median value and the absolute median deviation of 20 images, respectively.

With the aid of PentaMind, we further explored key drawing features associated with cognition including the shape and size of the pentagons. As anticipated, PentaMind assigned a higher cognition score to regular pentagons – a five-sided polygon in which all sides and angles are equal (Figure 5D). The proportionality of the size of the two pentagons was also correlated with a higher predicted cognitive score (Figure 5E). We observed a sharp decline in the prediction cognition when the size of the drawing was reduced (Figure 5F), which is in line with the fact that micrographia often seen in Parkinson’s disease^17^. Regarding line width, the cognition score decreased when the line was too thin but had a moderate effect otherwise (Figure 5G). Notably, the increase in line waviness had a strong influence on the prediction, even if the shape still appeared to be regular pentagons (Figure 5H). The line waviness likely reflects motor impairment, as drawing a straight line requires proper motor executions. Therefore, the model’s ability to quantify the line waviness may contribute to the enhanced accuracy of detecting motor-related cognitive impairment shown in Figure 3. This finding suggests line quality as a significant parameter for the cognitive assessment of hand drawing, which the conventional visual inspection mostly ignores.

## Discussion

This work revisited data from a traditional paper-and-pencil drawing test collected for decades with deep learning technology, bringing innovation to healthcare and life science. Results obtained with cutting-edge data analysis outperformed the standard human-based rating by about two-fold for predictions of cognitive function, revealed the novel relationships of hand-drawn pentagons with motor and related brain pathologies, and nominated specific drawing attributes affected by cognitive function. Our approach quantifies the contributions of diverse cognitive and motor neural systems underlying a commonly employed drawing task that can advance our understanding of the well-known association of cognitive and motor decline in older adults.

Our results suggest that quantifying facets of motor function underlying the drawn image was the primary source of the improved model performance compared to the conventional scoring. The subsequent analysis of brain pathologies, including cerebral atherosclerosis, arteriolosclerosis, and nigral neuronal loss also supported the involvement of motor function. For instance, cerebral atherosclerosis and arteriolosclerosis are associated with dementia and Parkinsonism^18^ by triggering neurovascular dysfunction such as decreased blood flow in the brain and impairment of blood-brain barrier integrity^19,20^. Also, the connection between the loss of nigral dopaminergic neurons and Parkinsonism is well-established^21^. Furthermore, the line waviness greatly impacted the predicted cognition, which also supports the connection between hand-motor execution and cognitive impairment. Recently, digital pen technology has enabled the measurement of intricate graphomotor data and shown its utility for cognitive evaluation^22,23^. Thus, incorporating time-series hand-drawing movement into the deep learning model may further improve the predictive accuracy for cognitive status. The success of this approach in highlighting the importance of quantifying sequential drawing for improved performance suggests that this approach may be useful for the assessment of other conventional motor skills, which currently assess only limited facets of the actual movement and do not even capture the duration of cognitive planning prior to the initial movement. Therefore, the significance of deep learning techniques will become even more crucial in analyzing other more complex behaviors, such as gait, whose 3D features are difficult to quantify during the routine clinical assessment of walking. These efforts may lead to a new lexicon of movement features derived from deep learning analysis that can be further examined using simulation, as was done in the current study.

We leveraged our cohorts’ racial diversity to examine the model’s applicability to white and African American populations. The model performed similarly for both races, indicating that the model is well-generalized across two races. This comparable performance may suggest that the number of training samples is sufficient to learn white and African-American-specific signals or that cognition-related drawing characteristics are independent of race. Clarifying these possibilities will guide the development of a generalized biomarker model based on hand drawings, which warrants further investigation. However, due to the small number of deceased people from the African American population (n=225), the findings on brain pathologies mostly reflect the data from the white population. Therefore, follow-up research is required to confirm the relationships between pentagon drawings and vascular pathologies in the African American population.

One of the challenges of using deep learning models is the need for more interpretability, or the ability to understand which image features are being used by the model to make predictions^24^. This is because deep learning models are often complex and have many layers, making it difficult to understand how the model reaches its conclusions. Various methods have been proposed to analyze an image’s most important parts for making predictions. These methods highlight the areas of an image that contain the most predictive information. Still, they may not explain which specific drawing characteristics are associated with cognitive decline. To address this challenge, we developed a fully parametrized simulator to generate a range of synthetic pentagon images. Then we analyzed the model’s predictions on these images to determine which features are most important for making a prediction. Having successfully identified a set of drawing attributes associated with predicted cognition, our approach could provide a novel way to explore handwritten biomarkers. Our result will serve as a resource to develop complementary computational approaches that automate the extraction of drawing features^25^. By quantifying each drawing attribute, we can examine the correlations between each attribute and the various phenotypes in greater depth.

Rapid and accurate assessment of complex behaviors resulting from diverse neural systems is critical to identifying the underlying neural mechanisms that can be targeted for treatment. By using deep learning technology to analyze handwriting images, it may be possible to design a faster and more accurate method for evaluating cognitive status. This could help healthcare providers more quickly identify individuals at risk for developing dementia and other cognitive impairments, allowing for early intervention and improved outcomes. Additionally, using handwriting samples as a biomarker for cognitive performance could provide insight into the molecular status of the brain, potentially advancing the development of precision medicine for dementia and other conditions. Overall, this study highlights the potential of using deep learning technology in healthcare to improve our understanding of cognitive impairments and to develop more effective treatment strategies.

## Method

### Study cohorts

All eligible participants were enrolled in one of three prospective aging studies at the Rush Alzheimer’s Disease Center (RADC), the Religious Orders Study (ROS)^26^, the Rush Memory and Aging Project (MAP)^26^, and the Minority Aging Research Study (MARS)^27^). These are prospective analytic community-based cohort studies. As community-based cohorts, the studies are far less susceptible to referral bias, which can introduce substantial sociodemographic, clinical, and genetic variations in patient research. At the time of enrollment, the average age was 77.4, the average length of education was 15.9 years, 72.9% were female, 77.4% were non-Latino white, and 21.9% were African American. All participants consent to undergo yearly comprehensive clinical examinations. Brain donation at the time of death is a condition of ROS and MAP study entry; it is optional for MARS. An Institutional Review Board of Rush University Medical Center approved all studies and participants gave written informed consent in accordance with the Declaration of Helsinki. As applicable, participants also sign an Anatomical Gift Act (AGA) for brain donation at death.

### Pentagon drawing test administration and pipeline processing

PDT was administered yearly to the participants as a part of items of the Mini-Mental State Examination (MMSE)^5^. The MMSE is a 30-item screener for gross cognitive impairment and dementia. It evaluates the severity of cognitive impairment across various cognitive domains. In one section, participants are asked to replicate a sample of intersecting pentagons on paper. The pentagons are then rated 0 (fail) or 1 (pass) based on the presence or absence of intersections and the number of vertices.

To prepare obtained pentagon drawing test data for training and testing the PentaMind model, we converted each pair of intersecting pentagons to a digital image. Then, we used an object detection method based on a C-Support Vector Classification algorithm (https://github.com/ttruty/object-detector) to identify and clip the region containing the intersecting pentagon. On the test set, the accuracy of the pentagon detection was 97.7%. In addition, we ran a manual evaluation to filter out images without pentagon drawings. The images were then padded with a white area around the edge of the pentagon to standardize the image size to 500 pixels by 500 pixels, while maintaining the size of the pentagon.

### Cognitive assessments

Each participant underwent comprehensive clinical evaluations at baseline and at each annual follow-up^28^. In summary, the cognitive battery includes 21 cognitive performance tests, 19 of which are used to develop a global composite measure of cognitive function (global cognition score) and 17 of which assess relatively dissociable cognitive domains, including episodic memory (7 measures), semantic memory (3 measures), working memory (3 measures), perceptual speed (2 measures), and visuospatial ability (2 measures).

### Motor assessments

Each participant was scored with two motor-related assessments: a global parkinsonism score and a global motor score. The global parkinsonism score was calculated by averaging the scores from each of the four parkinsonian domains^26^, which include bradykinesia, tremor, rigidity, and parkinsonian gait. The parkinsonian domains were assessed by qualified nurse clinicians using a modified version of the United Parkinson’s Disease Rating Scale (UPDRS)^29,30^. A higher score suggests more severe parkinsonian impairment of motor function. The global motor score is a summary of ten motor tests from four categories (1) hand strength (two items), (2) motor gait (four items), (3) motor dexterity (two items), and (4) motor balance (two items)^31^. To retain consistency with the global parkinsonism score, the sign of the global motor score and its components has been inverted such that a higher score reflects a more severe motor impairment. These two scores are independently linked with worse health outcomes when evaluated together^32^.

### Neuropathologic evaluations

We generated continuous measures for neuritic plaques, diffuse plaques, and neurofibrillary tangles. The modified NIA-Reagan criteria for diagnosing Alzheimer’s disease comprise the CERAD score for neuritic plaques and the Braak stage for neurofibrillary tangles^33^. Global AD pathology burden is a quantitative summary of AD pathology derived from counts of three AD pathologies: neuritic plaques, diffuse plaques, and neurofibrillary tangles, as determined by microscopic examination of silver-stained slides from 5 regions. For molecular specificity, we also quantified the load of parenchymal deposition of β-amyloid, and the density of abnormally phosphorylated paired helical filament tau (PHFtau)-positive neurofibrillary tangles, as previously described^34^. Additionally, we evaluated the extent of nigral neuronal loss, the presence of Lewy bodies^35^, the TDP-43 staging^36^, hippocampal sclerosis^37^, chronic macroscopic and microinfarcts^38^, cerebral amyloid angiopathy (CAA)^39^, the severity of atherosclerosis^40^, and arteriolosclerosis^40^.

### Model training

The convolutional neural network is a type of deep learning model that is particularly effective in image recognition tasks. Convolutional neural networks use a variation of the standard neural network architecture but with an added convolutional layer that helps to extract features from the input image. This convolutional layer comprises a set of filters, which are learned during the training process. These filters are used to scan over the input image, looking for meaningful patterns and features. The output of the convolutional layer is subsequently supplied to a series of fully connected layers, which perform the final classification or regression task. Convolutional neural networks excel at various tasks, including object detection, image classification, and image segmentation. The 47 vision recognition models implemented in Torchvision (v0.11.1) served as the foundation for the deep learning models that predict global cognition from pentagon drawings. Torchvision is a Pytorch-based library that provides several model architectures and pre-trained weights for computer vision. We modified the last layer of the model architectures such that the models output a single numeric number that corresponds to the global cognition score instead of image classes. The pre-trained weights against ImageNet^14^ were used to initialize the model parameters. The model parameters were then further tuned with a stochastic gradient descent (SDG) optimizer with a learning rate of 0.001, a momentum of 0.90, a weight decay of 0.0001, and a batch size of 32. In the case of AlexNet and SqueezeNet, a learning rate of 0.0005 was used. The input images were resized to 224 pixels by 224 pixels and augmented with imgaug (version 0.4.0)^41^. The maximum number of training epochs was set at 90, with an early termination threshold of 3 based on validation loss. The training pipeline was constructed with the workflow management system Snakemake^42^ and executed on the Google Cloud Platform with an NVIDIA Tesla T4 GPU.

### Pentagon simulation

We developed a simulator for drawing intersecting pentagons using the Matplotlib visualization library. In this simulation study, we examined eight drawing attributes, including angle distortion, line waviness, line width, the number of vertexes, alignment of two pentagons, the distance between pentagons, pentagon size, and size quality. For each parameter, we tested eight values that deviated from the standard pentagons. For each value of the main parameter tested, we generated 20 images with slight variations by randomly varying the rest of the parameters in a small range around the regular pentagons. A reproducible code and generated images are available at https://github.com/stasaki/. Generated images were supplied to the ten deep learning models, each of which had been trained with a distinct subset of data, and the predicted global cognition scores from the ten models were then averaged.

### Statistical analyses

To examine associations of the pentagon scores with clinical parameters and pathologies, we used Spearman’s correlation and linear regression as appropriate. For a multivariable linear regression model, the proportion of the variance explained by each variable was computed using the variance decomposition approach developed by Chevan and Sutherland^43^, implemented in the relaimpo R package^44^.

## Data availability

The data can be requested at the RADC Resource Sharing Hub at www.radc.rush.edu.

## Code availability

The trained model and the PDT simulator are available at https://github.com/stasaki/PentaMind/.

## Acknowledgments

We thank the study participants in ROS/MAP/MARS and the staff of the Rush Alzheimer’s Disease Center (RADC) and with support and data generation from NIH/NIA R01AG057911, R01AG061798, P30AG10161, P30AG72975, R01AG017917, R01AG015819, R01AG079133, R01AG22018, and R01AG042210.

## Author contributions

ST conceptualized and designed the study, constructed the predictive models, and performed the analysis. NK, TT, and ML prepared input pentagon drawing images. AZ contributed to the interpretation of the PentaMind model. ASB led the motor assessment. ST, NK, AZ, ASB, ML, and DAB interpreted the results. DAB secured funding, supervised resource allocation and data generation. ST drafted the manuscript, and all authors edited, read, and approved the submitted version.

