## Supplementary material for "Interpretable deep learning approach for extracting cognitive features from hand-drawn images of intersecting pentagons in older adults": Figure S1

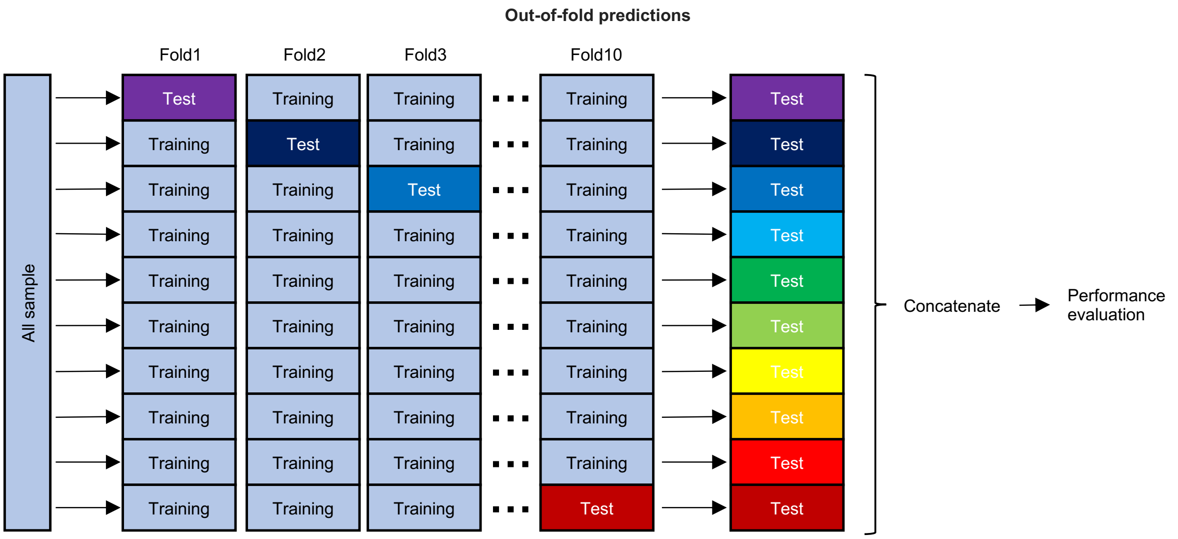


**Figure S1. The procedure of out-of-fold predictions.** We split 3,111 participants into 10 groups, selecting one group as the holdout test set. We then used data from the remaining groups to train PentaMind and generate a prediction for the test set. We repeated this process for all 10 folds, resulting in out-of-fold predictions for all 13,777 images from the 3,111 participants. Finally, we evaluated the performance of the model by comparing predicted cognition scores with actual values.
