## Supplementary material for "Interpretable deep learning approach for extracting cognitive features from hand-drawn images of intersecting pentagons in older adults": Figure S2

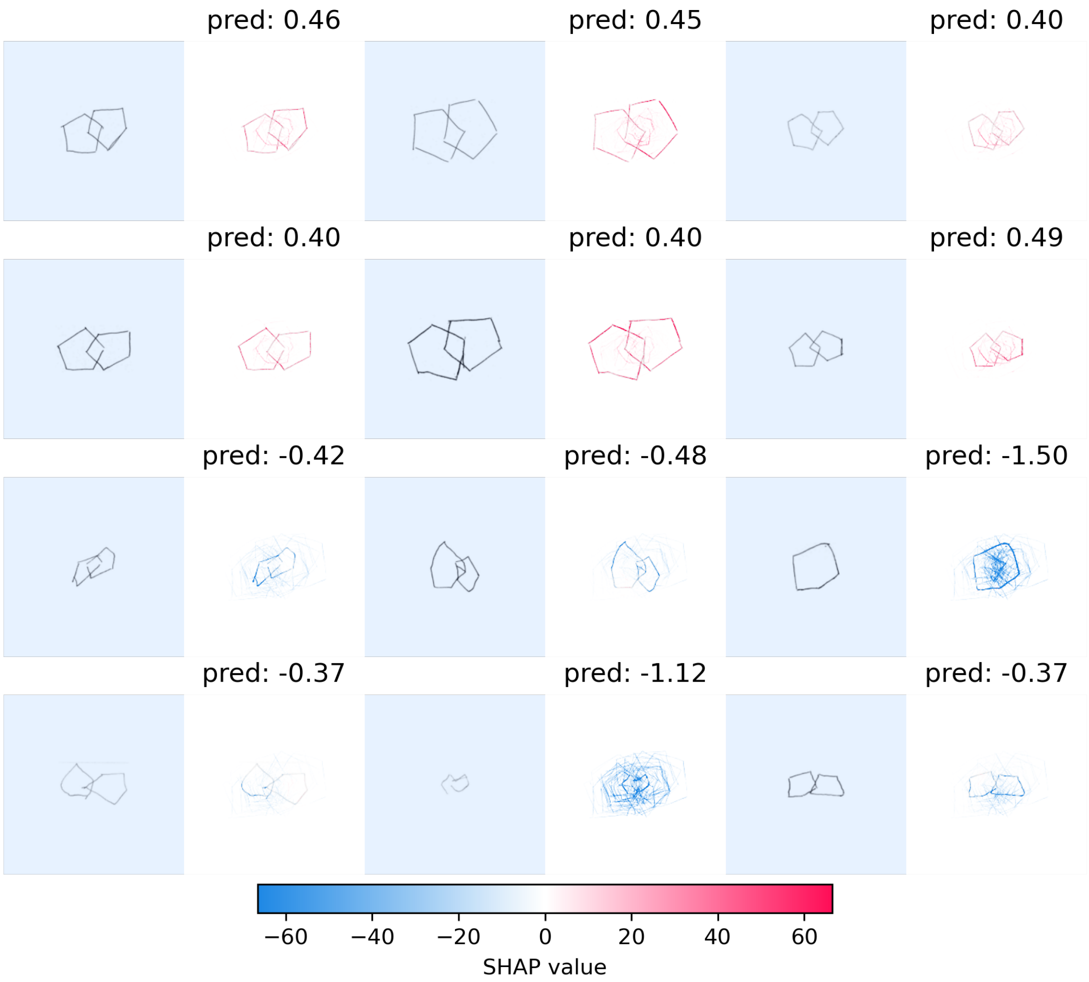


**Figure S2. DeepSHAP analysis that visualizes the elements influencing prediction.** We applied the DeepSHAP method to 6 pentagon drawings from non-cognitively impaired participants and 6 pentagon drawings from cognitively impaired participants. The red color indicates the parts that contribute to higher predicted cognition scores, while the blue color indicates the parts lowering the predicted cognition scores.
